# Structure-function analysis of histone H2B and PCNA ubiquitination dynamics using deubiquitinase-deficient strains

**DOI:** 10.1101/2023.06.18.545485

**Authors:** Kaitlin S. Radmall, Prakash K. Shukla, Andrew M. Leng, Mahesh B. Chandrasekharan

## Abstract

Post-translational covalent conjugation of ubiquitin onto proteins or ubiquitination is important in nearly all cellular processes. Steady-state ubiquitination of individual proteins *in vivo* is maintained by two countering enzymatic activities: conjugation of ubiquitin by E1, E2 and E3 enzymes and removal by deubiquitinases. Here, we deleted one or more genes encoding deubiquitinases in yeast and evaluated the requirements for ubiquitin conjugation onto a target protein. Our proof-of-principle studies demonstrate that absence of relevant deubiquitinase(s) provides a facile and versatile method that can be used to study the nuances of ubiquitin conjugation and deubiquitination of target proteins *in vivo*. We verified our method using mutants lacking the deubiquitinases Ubp8 and/or Ubp10 that remove ubiquitin from histone H2B or PCNA. Our studies reveal that the C-terminal coiled-domain of the adapter protein Lge1 and the C-terminal acidic tail of Rad6 E2 contribute to monoubiquitination of histone H2BK123, whereas the distal acidic residues of helix-4 of Rad6, but not the acidic tail, is required for monoubiquitination of PCNA. Further, charged substitution at alanine-120 in the H2B C-terminal helix adversely affected histone H2BK123 monoubiquitination by inhibiting Rad6-Bre1-mediated ubiquitin conjugation and by promoting Ubp8/Ubp10-mediated deubiquitination. In summary, absence of yeast deubiquitinases *UBP8* and/or *UBP10* allows uncovering the regulation of and requirements for ubiquitin addition and removal from their physiological substrates such as histone H2B or PCNA *in vivo*.

## Introduction

Ubiquitination (also called ubiquitylation) is the post-translational covalent conjugation of the 76 amino-acid ubiquitin onto proteins^1-3^. Ubiquitination is important in nearly all cellular processes^2^and is dysregulated many human diseases^4,5^. Protein ubiquitination is a reversible modification that is dynamically regulated by the actions of two opposing enzymatic activities^6^: ubiquitin conjugation and deubiquitination. E1 ubiquitin-activating enzymes, E2 ubiquitin-conjugating enzymes and E3 ubiquitin ligases ‘write’ or catalyze the addition of ubiquitin onto lysine residues in general in substrate proteins. Deubiquitinating enzymes (DUBs) ‘erase’ or remove the conjugated ubiquitin from target proteins. Thus, the *in vivo* steady state of ubiquitin conjugated onto any protein is a net result of these two counteracting enzymatic actions. We reasoned that deleting the genes coding for relevant DUB(s) could be used to evaluate ubiquitin conjugation step(s) onto a protein without complications from deubiquitination.

To establish this method, we focused on ubiquitination of two proteins, histone H2B and the PCNA, for which E2 and E3 enzymes and DUBs are well-characterized in budding yeast *Saccharomyces cerevisiae*. During transcription and other nuclear processes, the Rad6 E2 enzyme partners with a homodimer of the Bre1 E3 ligase and adapter protein Lge1 to conjugate a single ubiquitin onto histone H2B at lysine 123 (H2BK123ub1)^7-10^. This conjugated ubiquitin is targeted for removal by DUBs Ubp8 and Ubp10^11,12^. H2BK123ub1 controls nucleosome stability^13^ and chromatin dynamics^14^, and acts as a master ‘instructor’ modification to regulate the methylation of histone H3 at K4 and K79^15-18^. Following DNA damage, the Rad6-Rad18 E2-E3 complex catalyzes monoubiquitination of Pol30/PCNA (PCNAub1)^19,20^. This conjugated ubiquitin is removed by the DUB Ubp10^21^. To validate our *in vivo* ubiquitination assessment method, we created yeast deletion strains lacking *UBP8* and/or *UBP10*. Using the approach, we demonstrate that the C-terminal coiled-coil domain of Lge1 contributes significantly to Rad6-Bre1-mediated ubiquitin conjugation onto histone H2BK123. Like the coiled-coil domain of Lge1, the acidic tail of Rad6 is required for protein stability as well as its ubiquitin-conjugating activity onto H2BK123 *in vivo*. We also report that the distal acidic residues of helix-4 of Rad6 contribute to PCNA monoubiquitination. Parallel evaluation of H2BK123ub1 levels in the DUB deletion and wild-type strains revealed a role for the H2B C-terminal helix in regulating the dynamics of deubiquitination in addition to ubiquitin conjugation.

## Results

### Lge1 augments Rad6-Bre1-mediated histone H2BK123 monoubiquitination

In budding yeast, a complex comprised of Rad6, Bre1 and Lge1 catalyzes histone H2BK123ub1^7-9^, which in turn regulates histone H3K4 methylation^15,16,18,22^. H2BK123ub1 is strictly required for histone H3K4 trimethylation (me3) ^18^, a mark of active or activated state of transcription^23-26^. Matching well with these reported studies, immunoblots showed that H2BK123ub1 and H3K4me3 are abolished *in vivo* in yeast lacking Rad6 (*rad6Δ*) or Bre1 (*bre1Δ*) (Fig. 1A, compare lanes 1-3). Interestingly, H3K4 methylation was evident in the *lge1Δ* strain despite an apparent absence of H2BK123ub1 (Fig. 1A, compare lanes 1 and 4). This result suggested that H2BK123ub1 likely occurs in the absence of Lge1, and the seeming loss of this ubiquitination may be due to removal by DUBs Ubp8 and Ubp10. To test this possibility, we created a triple gene knockout yeast strain *ubp8Δubp10Δlge1Δ*, which lacks Lge1 and the DUBs Ubp8 and Ubp10. Additional triple gene knockout strains without Rad6 or without Bre1 and the two DUBs were also created as controls. Immunoblots showed that H2BK123ub1 was present in the *ubp8Δubp10Δlge1Δ* triple mutant but not in the control strains (Fig. 1B, compare lanes 2-4) or the *lge1Δ* mutant (Fig. 1A). The occurrence of H2BK123ub1 in the *ubp8Δubp10Δlge1Δ* mutant provided an explanation for the H3K4 methylation observed in the *lge1Δ* strain (Fig. 1A, lane 4). Collectively, these results confirmed that Rad6 and Bre1 catalyze monoubiquitination of histone H2BK123 in the absence of Lge1 and this modification is removed by DUBs Ubp8 and Ubp10.

**Figure 1.**
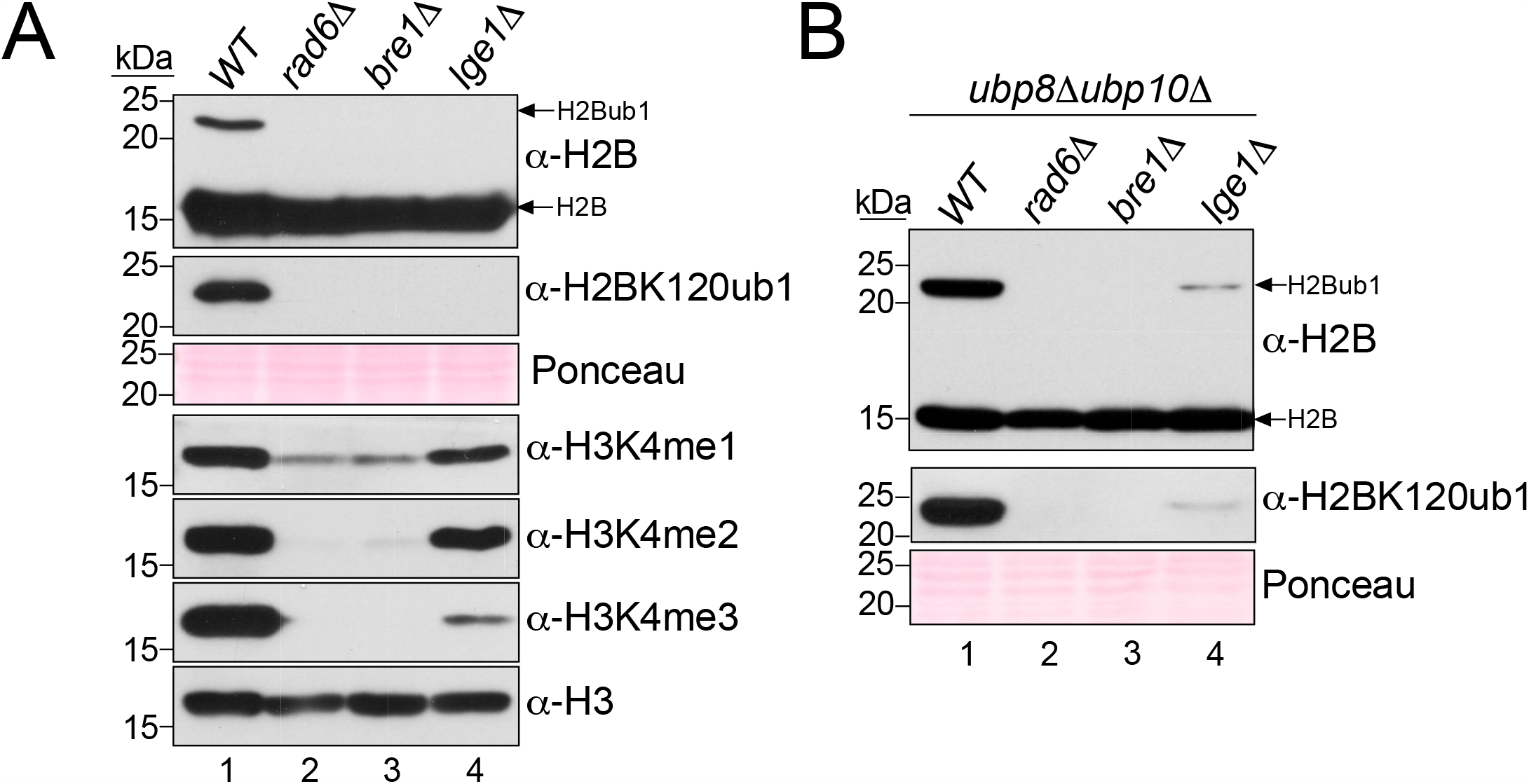
Low levels of H2BK123ub1 are detected in the absence of Lge1. **(A-B)** Immunoblots for histone H2BK123ub1 and/or histone H3K4 methylation (mono, me1; di, me2; tri, me3) in extracts prepared from A) strains lacking Rad6, Bre1 or Lge1 and B) strains lacking Rad6, Bre1 or Lge1 in the background of *ubp8Δubp10Δ* double deletion mutant. Ponceau S staining and histone H3 levels served as loading controls. Histone H2BK123ub1 was detected using anti-H2B or anti-H2BK120 ubiquityl antibody. Molecular weights of the protein standards used as size markers (kDa) are indicated.

Lge1 was proposed to act by blocking the action of DUB Ubp8^27^. However, H2BK123ub1 levels are reduced in the *ubp8Δubp10Δlge1Δ* triple mutant when compared to control *ubp8Δubp10Δ* double mutant that expresses wild-type (WT) Lge1 (Fig. 1B, compare lanes 1 and 4). This result indicated that Lge1 enhances Rad6-Bre1-mediated H2BK123 monoubiquitination *in vivo*. Overall, these results demonstrate a role for Lge1 in the ubiquitin ‘writing’ or conjugation step onto histone H2BK123 by Rad6-Bre1 E2-E3 enzymes. Further, these findings together provided the initial validation for use of yeast strains that lack DUBs as an experimental tool to evaluate protein ubiquitination *in vivo*.

### The C-terminal coiled-coil domain of Lge1 is critical for maintenance of histone H2BK123 monoubiquitination

Bre1 levels *in vivo* are dependent on Rad6^28,29^ and Lge1 levels are regulated by Bre1^30^. Consistent with these reports, analyses of mutants by immunoblotting showed that the steady-state Lge1 levels *in vivo* are dependent on Rad6 and Bre1 (Fig. S1A-B). We previously reported that an interaction with Rad6 stabilizes Bre1 *in vivo*^28^. A coiled-coil region in Bre1 is required for its interaction with and stabilization of Lge1^30^. However, the regions of Lge1 that are essential for maintenance of its steady-state levels and its functions in H2BK123ub1 were not known.

The structure of Lge1 predicted using AlphaFold2 contains an N-terminal unstructured region and a C-terminal helical coiled-coil domain^31,32^(Fig. 2A). To examine the contributions of these regions to Lge1 levels and functions *in vivo*, we created a series of N- or C-terminal truncation mutants of Lge1 (Fig. 2B) and expressed these proteins in the *lge1Δ* strain. Immunoblots showed that Lge1 levels were unaffected by the N-terminal truncations (Fig. 2C, Fig. S1C). In stark contrast, truncation of the C-terminus led to loss of Lge1 *in vivo* (Fig. 2C, compare lanes 6-9 to lane 2). Thus, the C-terminal coiled-coil region is essential for the steady-state levels of Lge1 *in vivo*. In *S. cerevisiae*, H2BK123ub1 via H3K4me3 regulates telomeric gene silencing^16,33,34^. Loss or reduction in these modifications leads to transcriptional activation of the silenced telomere-proximal *URA3* reporter, which compromises yeast growth on media containing counterselection agent 5-fluoroorotic acid (5FOA)^35^. The C-terminal truncation of Lge1 caused a severe telomeric silencing defect similar to that in the *lge1Δ* null mutant (Fig. 2C-D). The N-terminal truncation of residues 1-240 caused a subtle silencing defect when compared to the control strain expressing full-length Lge1 (Fig. 2D). Collectively, these silencing defects implicate both the N- and C-terminal regions of Lge1 in H2BK123ub1 formation.

**Figure 2.**
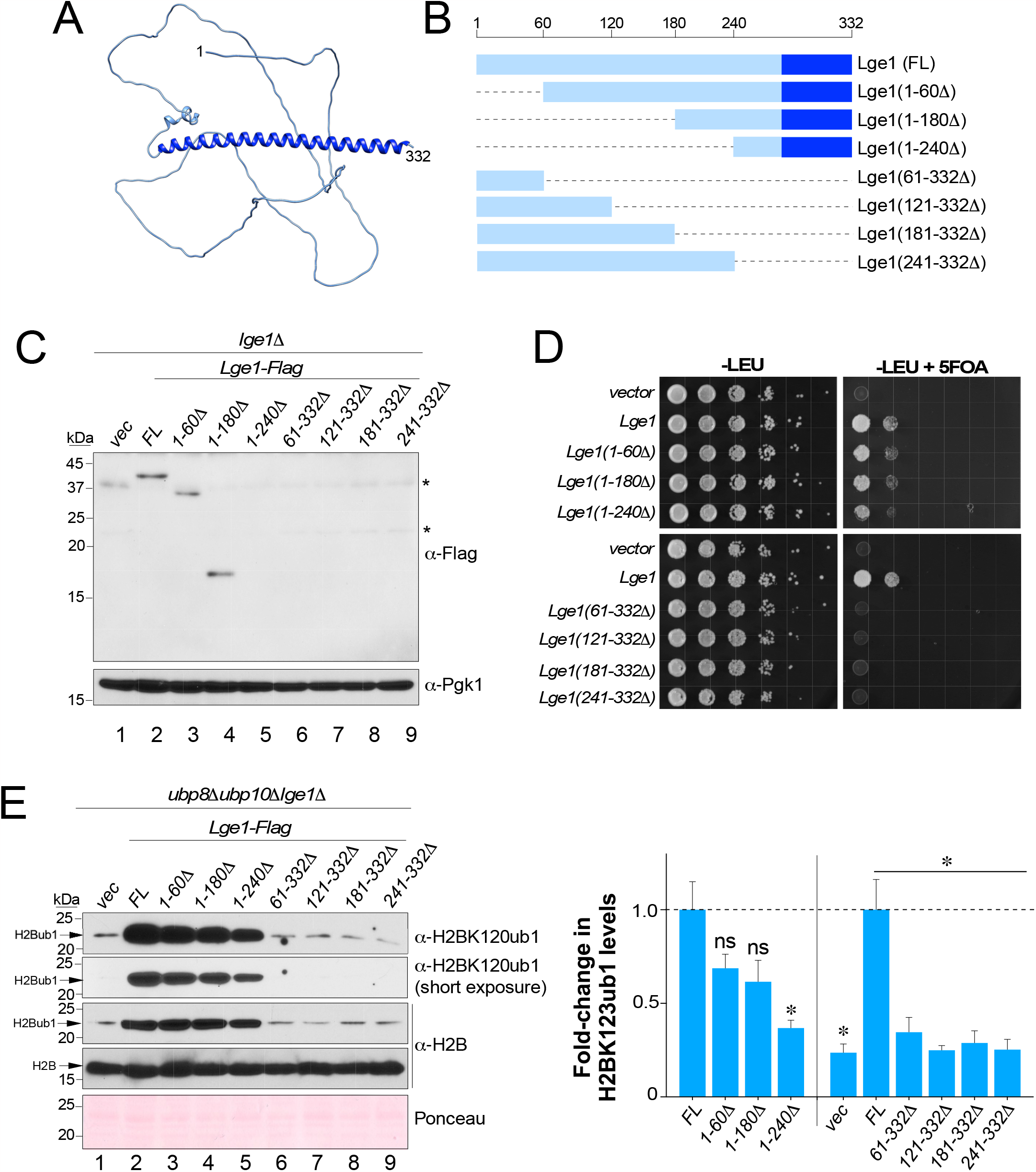
H2BK123ub1 and/or Lge1 levels *in vivo* are impaired by N- or C-terminal truncations of Lge1. **(A)** Structure of Lge1 predicted by AlphaFold2. **(B)** Schematic of N- and C-terminal truncation mutants of Lge1 compared to full-length (FL) Lge1. Dark blue box indicates coiled-coil domain. **(C)** Immunoblots for Flag epitope-tagged Lge1 or its truncation mutants expressed in *lge1Δ* null mutant strain. Extract from a strain transformed with empty vector (vec) served as negative control. Pgk1 levels served as loading control. Asterisk indicates cross-reacting proteins. **(D)** Growth assay for telomeric gene silencing was conducted by spotting 10-fold serial dilutions of indicated strains on synthetic medium lacking leucine (-LEU) or lacking leucine and containing 5-fluoroorotic acid (-LEU+FOA). **(E)** Left: Immunoblots for histone H2BK123ub1 in extracts prepared from *ubp8Δubp10Δlge1Δ* triple null-mutant transformed with either vector alone (vec) or a construct for Flag epitope-tagged full-length (FL) or indicated truncation mutants of Lge1. Ponceau S staining served as a loading control. Right: Fold-changes in H2BK123ub1 levels in the indicated truncation mutants relative to full-length Lge1 (set as 1). Right: plot of mean fold-change in H2BK123ub1 levels in the indicated mutants relative to full-length Lge1 (set as 1). Plotted are means ± SEM from three independent experiments. For densitometry quantitation, the signals for H2BK123ub1 in WT or mutant Lge1 were first normalized to the signals for Ponceau S-stained proteins, which served as loading control. ns, not significant; *\* p*-value <0.05 (Student’s t-test).

We used the DUB-deficient *in vivo* ubiquitination assessment approach to evaluate Lge1 function. We expressed the Lge1 deletion mutants in the *ubp8Δubp10Δlge1Δ* strain. Immunoblots showed that the expression of C-terminal truncation mutants that lack the coiled-coil domain all considerably decreased H2BK123ub1 levels similar to in the *lge1Δ* null mutant when compared to strain that expressed the full-length Lge1 (Fig. 2E, compare lanes 6-9 to lanes 1-2). H2BK123ub1 levels were also significantly reduced in cells that expressed Lge1 without N-terminal residues 1-240 (Fig. 2E, compare lanes 2-4 to lane 1). Overall, these results showed that the N-terminal IDR is largely dispensable, but the C-terminal coiled-coil domain is necessary for the function of Lge1 in Rad6-Bre1-mediated H2BK123ub1 formation *in vivo*.

### The acidic tail of Rad6 enhances Rad6-Bre1-mediated histone H2BK123 monoubiquitination

When compared to its homologs in *S. pombe* and *H. sapiens*, the *S. cerevisiae* Rad6 contains a 20 amino-acid C-terminal extension of mostly acidic residues (Fig. S2 and Fig. 3A-B). Unlike its structured globular UBC-fold domain (Fig. 3A), the acidic tail of Rad6 is protease-sensitive implying that it is disordered^36,37^. In *in vitro* assays, removal of the acidic tail of Rad6 severely inhibits mono- and poly-ubiquitination of histone H2B (Fig. 3C). Mutants of Rad6 that lack portions of the acidic tail, namely, *rad6-149* and *rad6-153*, were previously reported to impair *in vitro* activity^36,38,39^and *in vivo* functions such as sporulation and protein degradation^36,40^. However, the contributions of the Rad6 acidic tail to Rad6-Bre1-catalyzed monoubiquitination of histone H2BK123 and related *in vivo* functions remained unknown. In a telomeric silencing assay, the *rad6-149* mutant had a severe growth impairment on 5-fluoroorotic acid-containing medium or showed a silencing defect when compared to control strain that expresses wild-type (WT) Rad6 (Fig. 3D). In comparison, the *rad6-153* mutant or its derivatives had a subtle silencing defect on 5-fluoroorotic acid-containing medium compared to the control strain (Fig. 3D, compare third spots in the dilution series). Moreover, alanine substitution at all C-terminal non-alanine residues after position 149 caused a very severe silencing defect when compared to strains expressing WT Rad6 or other mutants (Fig. 3D). Collectively, these silencing defects suggested that the C-terminal acidic tail of Rad6 is important for H2BK123ub1 formation.

**Figure 3.**
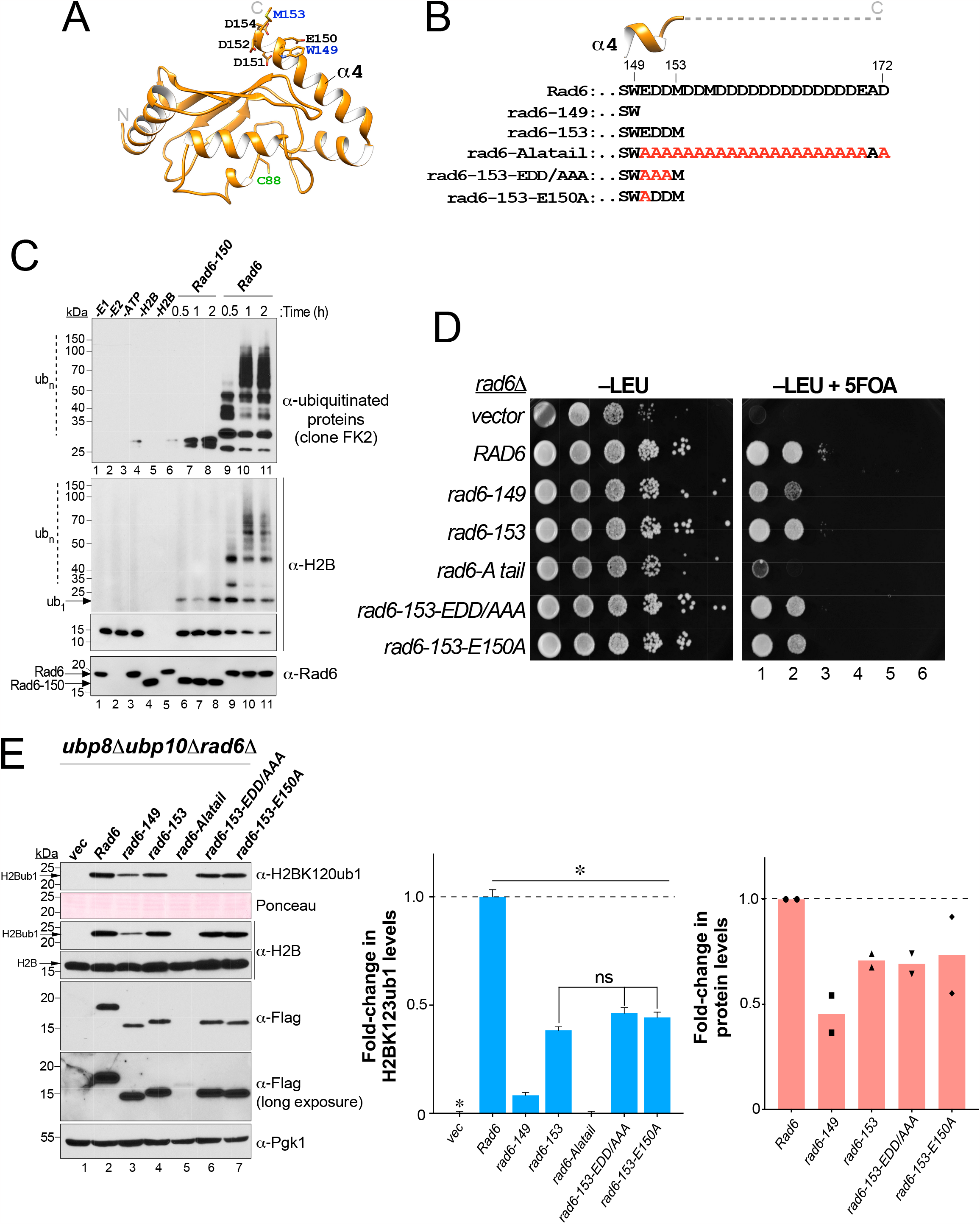
Deletion of C-terminal acidic tail decreases Rad6 and H2BK123ub1 levels *in vivo*. **(A)** Ribbon representation of Rad6 (PDB ID 1AYZ). Helix-4 and its terminal residues of Rad6 are indicated as is catalytic cysteine-88. **(B)** Sequences of the truncation and alanine substitution mutants are shown. **(C)** Immunoblots of products of an *in vitro* ubiquitination assay with recombinant WT Rad6 or Rad6-150 truncation mutant. Enzyme was incubated at 30°C for the indicated time along with the presence of ubiquitin (Ub), Uba1 (E1), ATP/Mg2+, and yeast histone H2B (substrate). The reaction mix was then resolved by SDS-PAGE prior to immunoblotting. ub_1_indicates monoubiquitinated H2B; ub_*n*_indicates polyubiquitinated substrate. **(D)** Growth assay for telomeric gene silencing conducted by spotting 10-fold serial dilutions of indicated strains on synthetic medium lacking leucine (-LEU) or lacking leucine and containing 5-fluoroorotic acid (-LEU+FOA). **(E)** Left: Immunoblots for histone H2BK123ub1 and either WT or mutant Rad6 in extracts prepared from *ubp8Δubp10Δrad6Δ* triple null-mutant transformed with either vector alone (vec) or constructs for Flag epitope-tagged full-length (FL) or the indicated truncation or point mutants of Rad6. Right: Fold-changes in H2BK123ub1 levels in the indicated mutants relative to full-length Rad6 (set as 1). Plotted are means ± SEM from three independent experiments. For densitometry quantitation, the signals for H2BK123ub1 in WT or mutant Rad6 were initially normalized to the signals for Ponceau S-stained proteins. ns, not significant; *\* p*-value <0.05 (Student’s t-test). Fold-changes in the indicated mutant Rad6 levels are shown relative to wild-type Rad6 (set as 1). Plotted are means from two independent experiments.

To test this possibility, we expressed either WT Rad6 or its C-terminal truncation or point mutants in the *ubp8Δubp10Δrad6Δ* strain. Immunoblots showed that H2BK123ub1 levels were significantly decreased in the strains expressing Rad6 mutants (*rad6-149* and *rad6-153*) that lacked the acidic tail when compared to the control strain that expresses full-length Rad6 (Fig. 3E, compare lanes 3-4 to lane 1). This result demonstrated the importance of the C-terminal acidic tail to Rad6’s histone H2B ubiquitination activity *in vivo*. Interestingly, expression of rad6-149 resulted in a larger decrease in H2BK123ub1 levels compared to the control than the expression of rad6-153 (Fig. 3E, compare lanes 3 and 4). This result indicated a functional role for residues 149-153 that are part of the conserved helix-4 of Rad6 (Fig. 3A-B). Structural studies showed that the distal residues in helix-4 of Rad6, especially glutamate 150 (E150), interact with Bre1^28^. Alanine substitutions at the three negatively charged residues (EDD/AAA) or at glutamate 150 (E150A) in the context of the rad6-153 sequence did not decrease H2BK123ub1 levels to the same extent as observed upon expression of rad6-149 (Fig. 3E, compare lanes 6 and 7 to lane 3). These results together suggested that an intact helix-4, but not its terminal charged residues, are required for Rad6-Bre1-mediated H2BK123ub1 formation.

It was reported that rad6-149 and rad6-153 are expressed at reduced levels relative to Rad6^36^. Indeed, immunoblots showed that the steady-state levels of rad6-149 and rad6-153 or its point mutant derivatives were reduced when compared to full-length Rad6 (Fig. 3E, compare lanes 3 and 4 to lane 2). The steady-state levels of Rad6 were nearly abolished by alanine substitution of all negatively charged residues in the tail (rad6-Alatail) (Fig. 3E, compare lane 5 to lane 2). The loss of Rad6 protein matches well with the absence of H2BK123ub1 and the severe silencing defect in this mutant (Fig. 3D-E). Overall, these results demonstrate that maintenance of steady-state levels of Rad6 and H2BK123ub1 *in vivo* depends on helix-4 and the acidic tail of Rad6.

### PCNA monoubiquitination depends on helix-4 of Rad6

Rad6 partners with Rad18 E3 ligase to monoubiquitinate Pol30 or PCNA (PCNAub1) following DNA damage^19^. The DUB Ubp10 removes the ubiquitin conjugated onto PCNA^21^. Hence, we deleted *UBP10* and *RAD6* in yeast (*ubp10Δrad6Δ*) to assess PCNAub1 formation. We used this method to investigate the effects of C-terminal truncation or point mutations of Rad6 on PCNAub1 formation. Consistent with the low levels of rad6-Alatail (Fig. 3E, lane 5), PCNAub1 formation was nearly abolished in the rad6-Alatail mutant compared to control cells that express WT Rad6 (Fig. 4A, compare lane 5 to lane 2). The steady-state levels of PCNAub1 were not altered upon expression of rad6-153 but were significantly decreased in the presence of rad6-149 when compared to the control strain with full-length Rad6 (Fig. 4A-B). This finding matches well with the previous report that *rad6-149*, but not *rad6-153*, is UV sensitive^36^. PCNAub1 levels were not reduced in the rad6-153 mutant, but mutating the distal charged residues in helix-4 (rad6-153-EDD/AAA) significantly reduced PCNAub1 levels when compared to rad6-153 or wild-type Rad6 (Fig. 4A-B, compare lane 6 to lanes 2 and 4), and similar to those in the cells that express rad6-149, which lack these residues (Fig. 4A-B, compare lane 6 to lanes 2-3). Thus, the acidic tail of Rad6 is dispensable, but the C-terminal negatively charged residues of helix-4 are vital for Rad6-Rad18 catalyzed PCNAub1 formation.

**Figure 4.**
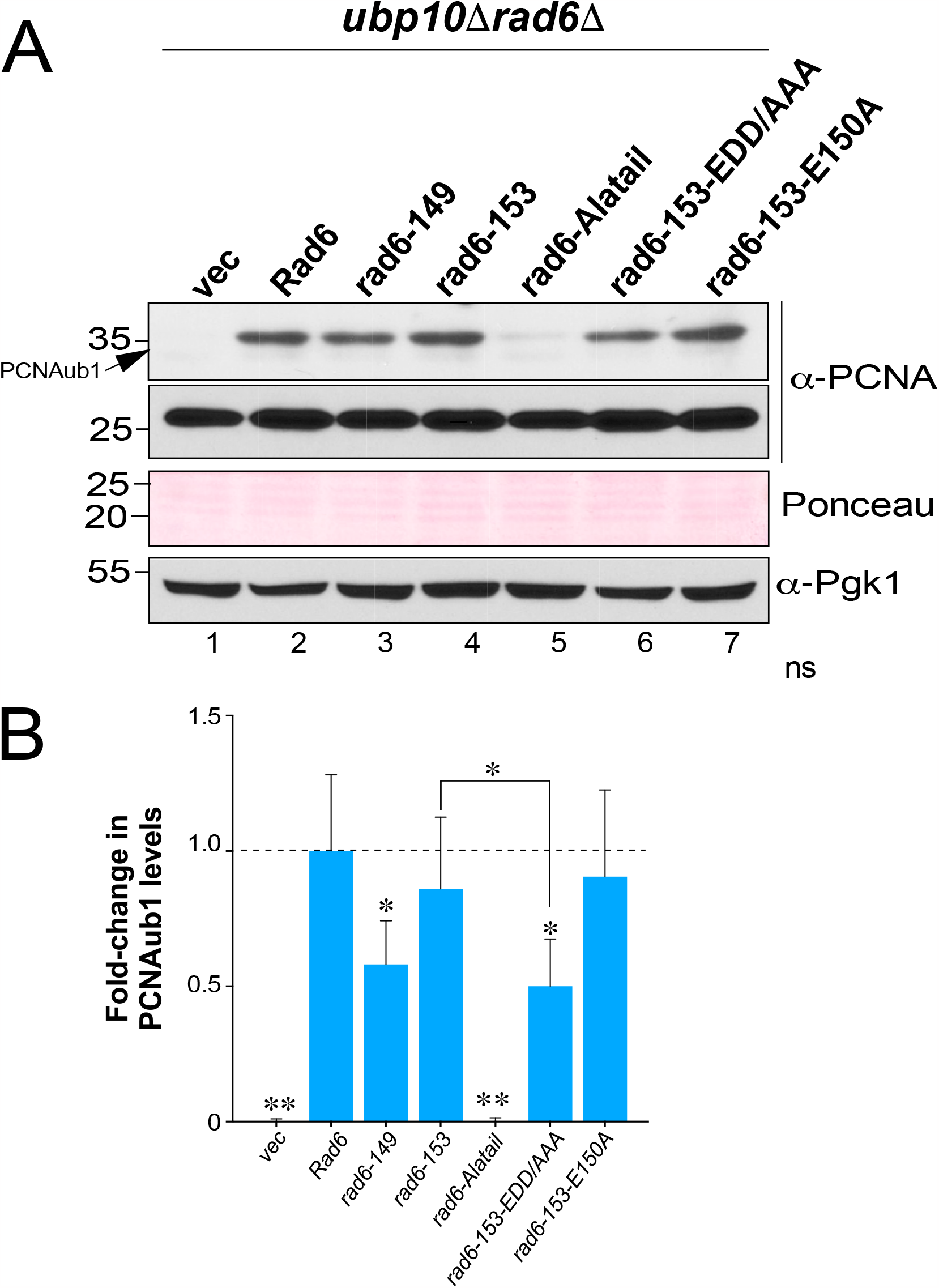
Mutations in helix-4 terminus impair PCNA ubiquitination *in vivo*. **(A)** Immunoblots for PCNAub1 in extracts prepared from *ubp10Δrad6Δ* null-mutant transformed with either vector alone (vec) or constructs for Flag epitope-tagged full-length Rad6 or its indicated truncation or point mutants. Ponceau S staining and Pgk1 levels served as loading controls. **(B)** Fold-change in PCNAub1 levels in the indicated mutants relative to full-length Rad6 (set as 1). For densitometry quantitation, the signals for PCNAub1 in WT or mutant Rad6 were initially normalized to the signals for Ponceau S-stained proteins. Plotted are means ± SEM from three independent experiments. *\*, p*-value <0.05; *\*\*, p*-value <0.001 (Student’s t-test).

### Histone H2B alanine-120 mutations influence ubiquitin conjugation and deubiquitination steps

We previously reported that mutations in arginine 119 and threonine 122 in the histone H2B C-terminal helix altered the levels of monoubiquitination at lysine 123 (K123)^34^. In our structure-function studies of the H2B C-terminal helix (Cα), we created yeast strains that expressed charged aspartate (D) or arginine (R) substitution at alanine 120 (A120) (Fig. 5A). Both *H2B-A120D* and *H2B-A120R* mutants showed a severe telomeric silencing defect (Fig. 5B), suggesting that these histone H2B mutations adversely affected H2BK123 monoubiquitination. Indeed, immunoblotting showed that the steady-state levels of H2BK123ub1 were significantly reduced in *H2B-A120D* and *H2B-A120R* mutants when compared to control strain expressing WT H2B (Fig. 5C). Next, we expressed either WT or the A120 mutant as the sole type of histone H2B in an *ubp8Δubp10Δ*-based histone shuffle strain. Immunoblots showed that H2BK123ub1 levels were significantly decreased in the H2B-A120D mutant compared to control with WT H2B even in the absence of DUBs (Fig. 5D), which indicated that aspartate substitution at residue 120 of histone H2B inhibits the Rad6-Bre1-Lge1-catalyzed H2BK123 monoubiquitination. H2BK123ub1 levels in the H2B-A120R mutant were similar to those in the DUB null mutant *ubp8Δubp10Δ* strain that expressed WT H2B (Fig. 5D). This is in contrast to the decreased H2BK123ub1 levels when the H2BA120R mutant was expressed in a strain that contained DUBs Ubp8 and Ubp10 (*UBP8UBP10*) (Fig. 5C). These results revealed that decreased H2BK123ub1 levels in the presence of H2B-A120R mutant in the strain expressing DUBs is not due to inhibition of Rad6-Bre1-Lge1 catalyzed ubiquitin conjugation but is instead due to enhanced removal of the conjugated ubiquitin by Ubp8 and Ubp10. Taken together, these findings demonstrate that the DUB deficiency provides an *in vivo* ubiquitination assessment method that can yield insights into the process of deubiquitination in addition to ubiquitin conjugation.

**Figure 5.**
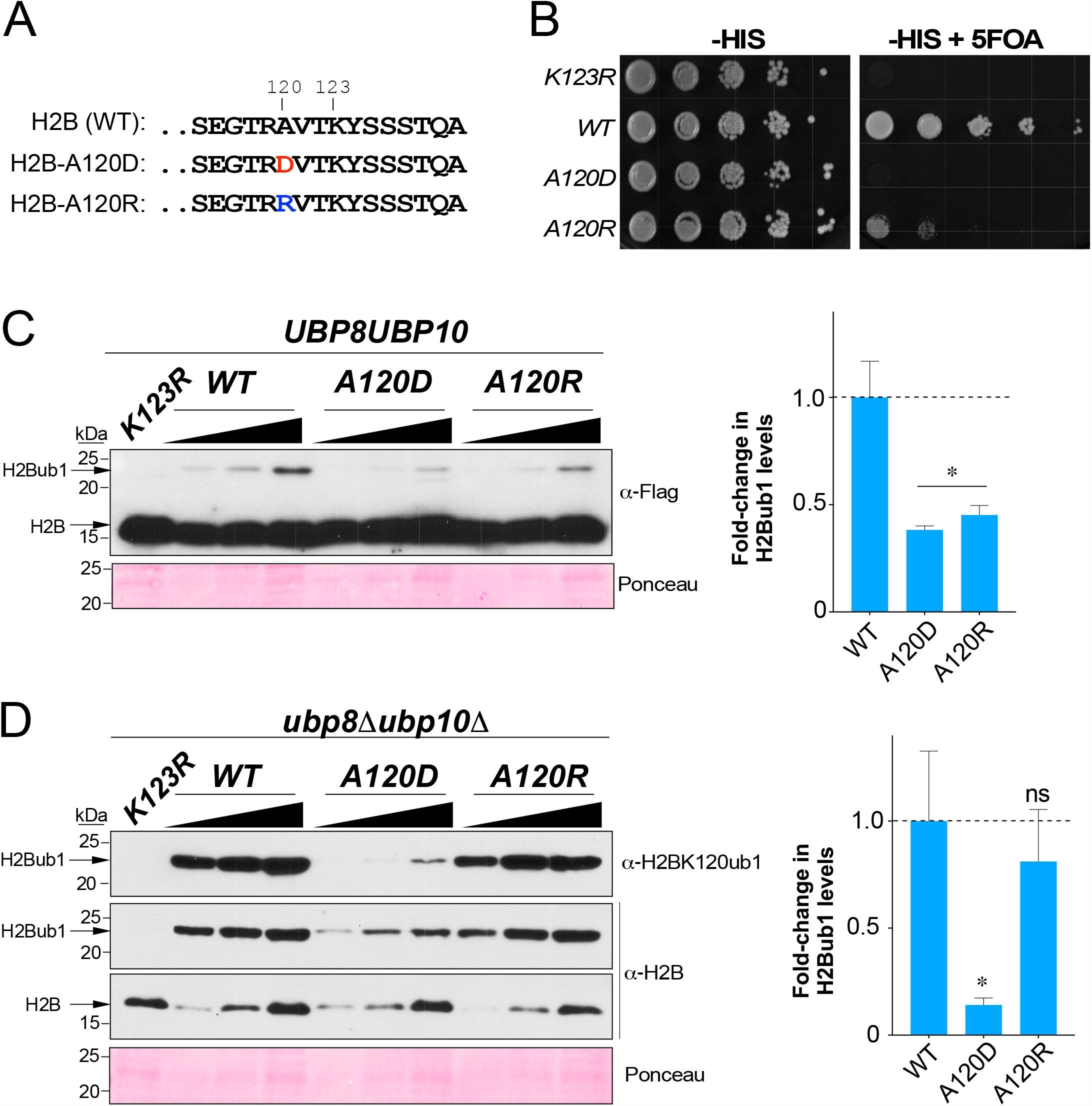
Substitution of alanine 120 in the H2B C-terminal helix with charged amino acid interferes with ubiquitination at lysine 123. **(A)** Sequences of the distal end of WT H2B C-terminal helix and of mutants. Lysine 123, the site of monoubiquitination is indicated. **(B)** Growth assay conducted by spotting 10-fold serial dilutions of indicated strains on synthetic medium lacking histidine (-HIS) or lacking histidine and containing 5-fluoroorotic acid (-HIS+FOA). **(C-D)** Left: Immunoblots for H2BK123ub1 in extracts prepared from C) *UBP8UBP10* or D) *ubp8Δubp10Δ* strains expressing Flag epitope-tagged WT or mutant histone H2B. *Triangles* denote increasing amounts of extracts used. Ponceau S staining served as loading control. Right: Fold-change in H2BK123ub1 levels in the indicated mutants relative to WT H2B (set as 1). For densitometry quantitation, the signals for H2BK123ub1 in WT or mutant H2B were initially normalized to the signals for Ponceau S-stained proteins. Plotted are means ± SEM from three independent experiments. ns, not significant; *\*, p*-value <0.05 (Student’s t-test).

## Discussion

In this study, we report a method in *S. cerevisiae* that allows study of the dynamics of ubiquitination of a substrate or target protein *in vivo*. This approach is applicable when the regulatory enzymes or factors involved in ‘writing’ or ‘erasing’ the ubiquitin modification from a target protein are known (Fig. 6A). This is the case for histone H2BK123 monoubiquitination. In yeast, approximately 10% of histone H2B is monoubiquitinated at steady-state; this level is maintained by the actions of ‘writer’ Rad6-Bre1-Lge1 complex^7-9,27^and ‘eraser’ deubiquitinases (DUBs) Ubp8 and Ubp10^8,11,12^(Fig. 6B). In the absence of Ubp8- and Ubp10-mediated removal, ubiquitination of histone H2BK123 catalyzed by the Rad6-Bre1-Lge1 complex reaches very high levels (>6-12-fold in *ubp8Δubp10Δ* relative to *UBP8UBP10*)^12,41^(Fig. 6B-C). However, loss of Lge1 regions or the C-terminal acidic tail of Rad6 impairs the production of high levels of H2BK123 monoubiquitination despite the absence of Ubp8 and Ubp10 (Fig. 6D). Thus, experiments performed using our *in vivo* ubiquitination assessment method revealed that Lge1 and the acidic tail of Rad6 are required for the monoubiquitination of histone H2B.

**Figure 6.**
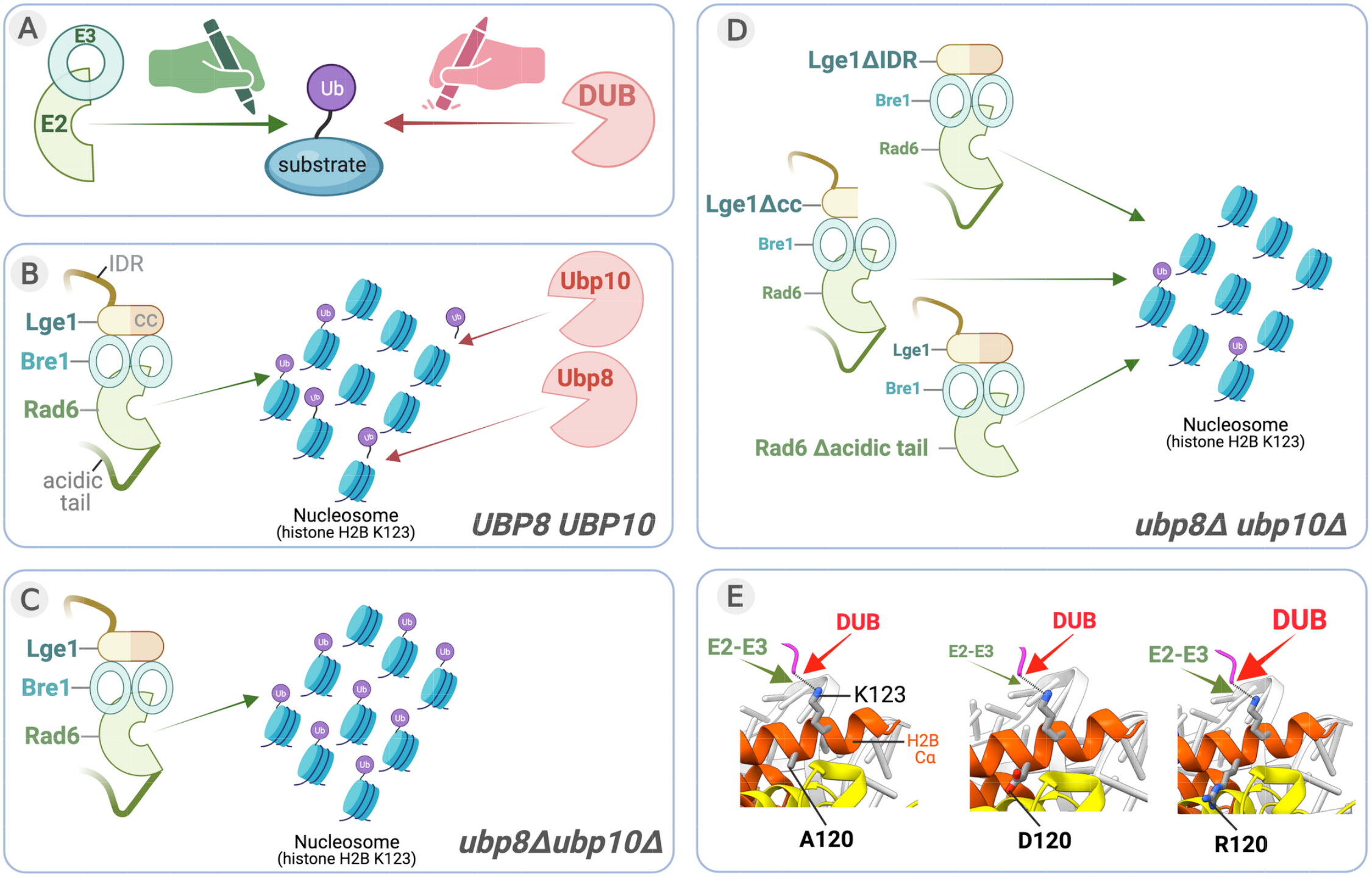
An *in vivo* DUB deletion mutant-based approach for evaluating ubiquitin conjugation and deubiquitination. **(A)** The steady-state levels of ubiquitination of a protein *in vivo* are maintained by the actions of ‘writer’ E2 ubiquitin-conjugating enzymes and E3 ubiquitin ligases and ‘eraser’ DUBs. **(B)** In yeast *S. cerevisiae*, the *in vivo* steady-state levels of H2BK123ub1 in a nucleosome are maintained by the ‘writer’ complex comprised of Rad6 (E2), Bre1 (E3) and accessory/adapter protein Lge1, and two ‘eraser’ DUBs, Ubp10 and the SAGA complex-associated Ubp8. IDR, intrinsically disordered region; cc, coiled-coil domain. **(C)** In the *ubp8Δubp10Δ* double null mutant strain, H2BK123ub1 accumulates due to ubiquitin addition and absence of deubiquitination. **(D)** High levels of H2BK123ub1 are not observed in the *ubp8Δubp10Δ* mutant strain when either the IDR or coiled-coil domain of Lge1 or the C-terminal acidic tail of Rad6 are deleted. Thus, the absence of relevant DUBs revealed the roles for various regions or domains of proteins involved in the ubiquitin-conjugation step. **(E)** Residues of the H2B C-terminal helix (Cα) impact the activity of the E2-E3 complex and the DUBs by influencing their access to substrate K123 or its ubiquitin conjugated form, respectively. Aspartate substitution at residue 120 in H2B Cα inhibits the Rad6-Bre1-Lge1-mediated monoubiquitination of H2BK123 in both WT and *ubp8Δubp10Δ* strains. In contrast, arginine substitution at position 120 in H2B Cα promotes removal of the conjugated ubiquitin by Ubp8 and Ubp10, as evidenced by the reduced H2BK123ub1 in the *H2BA120R* mutation in a strain expressing these two DUBs and not in their absence. Thus, the use of the DUB deletion strain informed on the dynamics of deubiquitination in addition to the ubiquitin conjugation.

Bre1 levels *in vivo* are dependent on Rad6 and its interaction with Rad6^28^. We showed that Lge1 levels *in vivo* are dependent on Rad6 and Bre1 (Fig. S1A-B). The C-terminal coiled-coil domain of Lge1 binds Bre1^27,42^, and our data indicated that this domain is essential for maintenance of Lge1 levels *in vivo* (Fig. 2C). Thus, our findings, along with others, indicate to a hierarchical relationship between the subunits of the yeast histone H2B ubiquitin-conjugating complex: Rad6, the E2 ubiquitin conjugase, forms the core of the complex and interacts with and stabilizes the homodimer of the E3 ligase Bre1, which in turn binds and stabilizes Lge1 via interaction with the C-terminal coil domain.

Intrinsically disordered regions (IDRs) in chromatin or transcriptional regulators can act as degrons or in degron masking to influence protein stability^43^. However, loss of the N-terminal IDR of Lge1 did not affect its stability *in vivo*. Expression of mutant Lge1 lacking the IDR decreased H2BK123ub1 levels within gene bodies^42^, which indicates a role for the IDR in enhancing Rad6-Bre1-mediated H2BK123ub1 formation during transcription elongation. IDRs also mediate protein-protein interactions to regulate protein activity^43^. Lge1 interacts with multiple proteins in addition to Bre1 *in vivo*^44^. Using our *in vivo* ubiquitination assessment method, we showed that the IDR of Lge1 contributes to the ubiquitin conjugation onto histone H2BK123 catalyzed by the Rad6-Bre1 E2:E3 enzymes. It is possible that the IDR of Lge1 stimulates Rad6-Bre1-mediated conjugation of ubiquitin onto H2BK123 through an allosteric mechanism via its transient interactions with various factors involved in transcription elongation.

The C-terminal acidic tail of Rad6 forms a disordered region^36,37^and is important for *in vitro* activity^38^and certain *in vivo* functions^39,40^. The C-terminal acidic tail of the E2 enzyme Cdc34 mediates its interaction with SCF^Cdc4^ E3 ligase^45^. However, the Bre1 E3 ligase binds Rad6 even in the absence of Rad6’s C-terminal acidic tail^28^. We discovered that the presence of acidic tail stimulates Rad6 activity *in vitro* in the absence of an E3 ligase. Here we showed that the C-terminal acidic tail contributes to protein stability *in vivo* and that charge-neutralizing alanine substitutions caused a near complete elimination of Rad6 from the yeast cell. Overall, our observations from the *in vivo* ubiquitination assessment method revealed that the acidic tail of Rad6 contributes to the Rad6-Bre1-mediated ubiquitin conjugation onto histone H2BK123. We speculate that the C-terminal acidic tail stabilizes an enzymatically active conformation of Rad6.

Helix-4 is a conserved integral constituent of the E2 UBC fold^46-48^. Structural studies have shown that E150 in helix-4 of Rad6 contacts Bre1 K31^28^. Our studies showed that an intact helix-4, but not its last three negatively charged residues, is required for normal levels of Rad6 and H2BK123ub1. Superposition of the crystal structures of yeast Rad6 onto human Rad6b/UBE2B in complex with the Rad6-binding domain of Rad18 shows that D151 in helix-4 of Rad6 can potentially interact with Rad18 (Fig. S3). In support of this, we found that the distal EDD residues of helix-4 are vital for efficient monoubiquitination of PCNA by the Rad6-Rad18 E2-E3 complex.

Residues in the H2B C-terminal helix (Cα) alter the dynamics of ubiquitination and deubiquitination at K123 by affecting chromatin binding and/or activities of Rad6-Bre1 E2-E3 or DUBs Ubp8 and Ubp10, as shown previously^34^. Using our ubiquitination assessment method, we established that charged substitutions at H2B Cα A120 differentially impact the ubiquitin addition or removal processes at K123 (Fig. 6E). Structure-based modeling showed that a negatively charged aspartate or a positively charged arginine substitution at A120 would result in repulsion or attraction, respectively, due to their presence in a primarily acidic neighborhood in the nucleosome (Fig. S4). In turn, these electrostatic forces could change the conformation of the C-terminal helix of H2B to alter substrate accessibility such that an aspartate substitution at position 120 inhibits Rad6-Bre1-mediated ubiquitin conjugation onto K123, whereas an arginine substitution at this position promotes Ubp8 and/or Ubp10-mediated H2B K123 deubiquitination. In sum, this example along with others demonstrate that the use of DUB-deletion strains provides a versatile approach to reveal the nuances and dynamics of ubiquitin addition as well removal from a target protein.

## Methods

### Plasmid construction

To create Lge1 expression constructs, the *LGE1* promoter fragment (490 bp) upstream of the start codon was PCR amplified to contain Not1 and BamH1 sites at 5’ and 3’ end, respectively. The *LGE1* terminator region (499 bp) downstream of the stop codon was PCR amplified to contain the sequence coding for the FLAG epitope and additionally contain Spe1 and Xho1 restriction sites at 5’ and 3’ ends, respectively. The *LGE1* promoter and terminator amplicons were then inserted into Not1-Xho1-digested vector pRS304 (*TRP1, CEN*)^49^using NEBuilder® HiFi DNA Assembly (NEB) to obtain pMC12. The full-length *LGE1* coding region and its truncation mutants were PCR amplified from genomic DNA isolated from the parental strain DHY217 as the template and inserted into BamH1-Spe1-digested pMC12. The *LGE1prom-LGE1 (WT or mutant)-Flag-LGE1term* fragment in pRS304 backbone was digested with Not1 and Xho1 and inserted into the same sites in pRS305 (*LEU2, CEN*)^49^. These pRS305-based constructs were linearized with Spe1 and a PCR product containing the coding sequence for 8 copies of the V5 epitope tag, and a stop codon was then inserted by sequence and ligation independent cloning (SLIC)^50^.

To create Rad6 expression constructs, the *RAD6promoter-RAD6-Flag-RAD6terminator* expression cassette was excised as a Kpn1-Sac1 fragment from construct pMC7^28^ and inserted into the same sites in vector pRS41H^51^. This construct was digested with Spe1 and BamH1 and either truncation or point mutants of Rad6 were generated by PCR amplification or by synthesis of Integrated DNA Technologies (IDT) gblocks® gene fragment, which were then inserted using NEBuilder® HiFi DNA Assembly kit (NEB). The expression cassette for WT or mutant *RAD6* in the pRS41H backbone was excised with Kpn1 and Sac1 and inserted into the same sites in vector pRS305 (*LEU2, CEN*). Aspartate or arginine substitution mutations at residue 120 of histone H2B were introduced by PCR-based site-directed mutagenesis in pZS145 (*HTA1-Flag-HTB1 CEN HIS3*)^16^. For bacterial expression, the codon optimized coding sequence for full-length Rad6 or its truncation mutant lacking the acidic tail (Rad6-150) were cloned into pET28a. All plasmid constructs were confirmed by Sanger or Nanopore sequencing.

### Yeast strains and media

Yeast cells were grown in YPAD broth (1% yeast extract, 2% peptone, 2% dextrose, and 0.004% adenine hemisulfate) or in synthetic dropout (SD) media. To prepare solid media, 2% agar was added to liquid broth prior to autoclaving. Gene knockout strains used in this study were created in parental YMH171 and/or DHY214/DHY217 strains reported previously^52,53^. Briefly, to create a gene knockout strain, the coding region was replaced with either an antibiotic resistance or auxotrophic marker gene using PCR products amplified from genomic DNA isolated from the respective deletion mutant obtained from a commercial source or using the strain available in our lab collection or, alternatively, using pF6a-KanMX or pAG25 ^54^or the relevant pYM series vector ^55^as the template. *RAD6*-null mutant strains were also created using a construct containing *URA3* in place of the *RAD6* coding region flanked by *RAD6* promoter and terminator sequences, which was linearized with HindIII-BamH1 prior to transformation. H2B mutant strains were created using plasmid shuffle approach in parental strain Y131^8^. Genotypes of yeast strains are listed in Table 1.

**Table 1.**
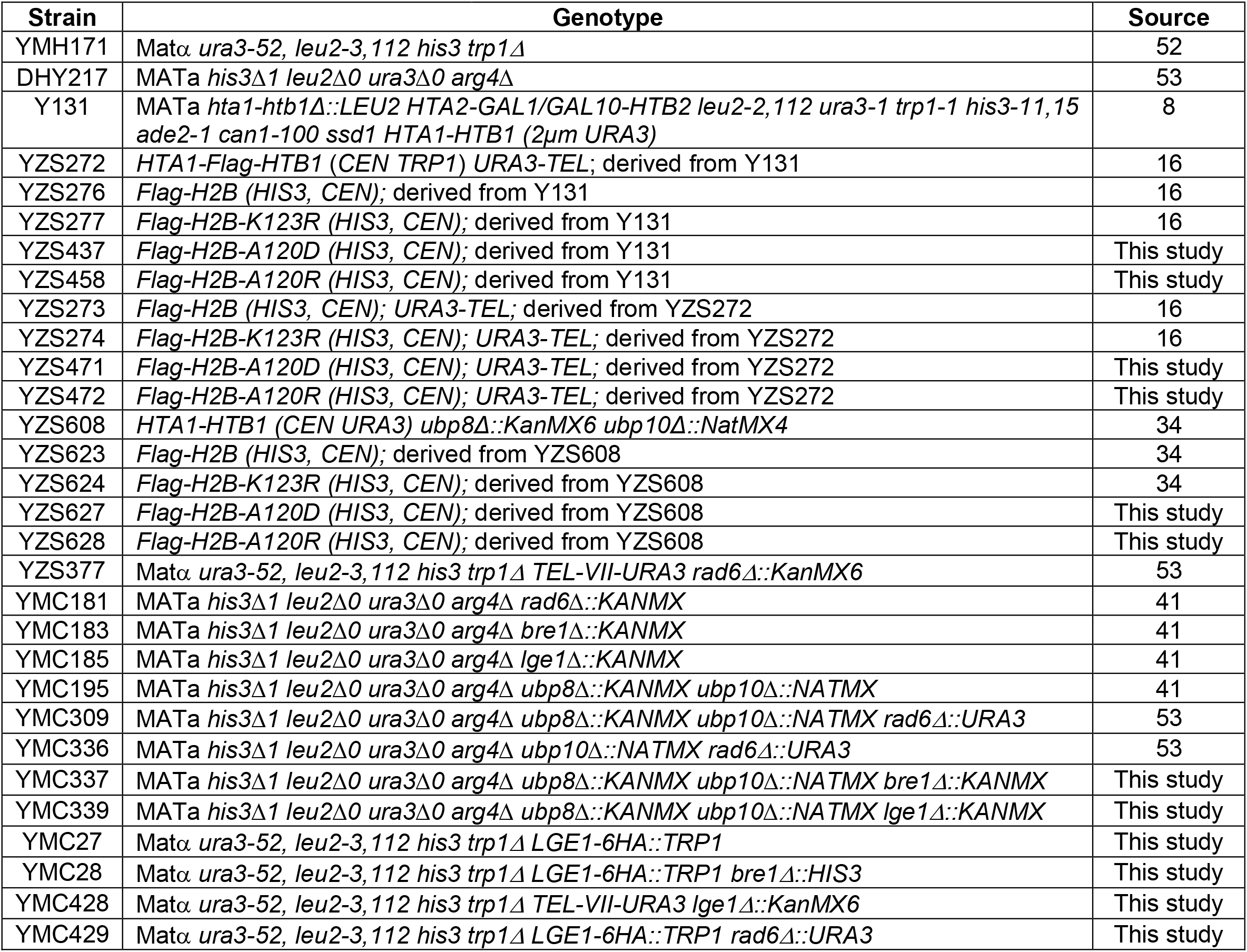
Yeast strains used in this study.

| **Strain** | **Genotype** | **Source** |
| --- | --- | --- |
| YMH171 | *Matα ura3-52, leu2-3,112 his3 trp1Δ* | 52 |
| DHY217 | *MATa his3Δ1 leu2Δ0 ura3Δ0 arg4Δ* | 53 |
| Y131 | *MATa hta1-htb1Δ::LEU2 HTA2-GAL1/GAL10-HTB2 leu2-2,112 ura3-1 trp1-1 his3-11,15 ade2-1 can1-100 ssd1 HTA1-HTB1 (2μm URA3)* | 8 |
| YZS272 | *HTA1-Flag-HTB1 (CEN TRP1) URA3-TEL*; derived from Y131 | 16 |
| YZS276 | *Flag-H2B (HIS3, CEN)*; derived from Y131 | 16 |
| YZS277 | *Flag-H2B-K123R (HIS3, CEN)*; derived from Y131 | 16 |
| YZS437 | *Flag-H2B-A120D (HIS3, CEN)*; derived from Y131 | This study |
| YZS458 | *Flag-H2B-A120R (HIS3, CEN)*; derived from Y131 | This study |
| YZS273 | *Flag-H2B (HIS3, CEN); URA3-TEL*; derived from YZS272 | 16 |
| YZS274 | *Flag-H2B-K123R (HIS3, CEN); URA3-TEL*; derived from YZS272 | 16 |
| YZS471 | *Flag-H2B-A120D (HIS3, CEN); URA3-TEL*; derived from YZS272 | This study |
| YZS472 | *Flag-H2B-A120R (HIS3, CEN); URA3-TEL*; derived from YZS272 | This study |
| YZS608 | *HTA1-HTB1 (CEN URA3) ubp8Δ::KanMX6 ubp10Δ::NatMX4* | 34 |
| YZS623 | *Flag-H2B (HIS3, CEN)*; derived from YZS608 | 34 |
| YZS624 | *Flag-H2B-K123R (HIS3, CEN)*; derived from YZS608 | 34 |
| YZS627 | *Flag-H2B-A120D (HIS3, CEN)*; derived from YZS608 | This study |
| YZS628 | *Flag-H2B-A120R (HIS3, CEN)*; derived from YZS608 | This study |
| YZS377 | *Matα ura3-52, leu2-3,112 his3 trp1Δ TEL-VII-URA3 rad6Δ::KanMX6* | 53 |
| YMC181 | *MATa his3Δ1 leu2Δ0 ura3Δ0 arg4Δ rad6Δ::KANMX* | 41 |
| YMC183 | *MATa his3Δ1 leu2Δ0 ura3Δ0 arg4Δ bre1Δ::KANMX* | 41 |
| YMC185 | *MATa his3Δ1 leu2Δ0 ura3Δ0 arg4Δ lge1Δ::KANMX* | 41 |
| YMC195 | *MATa his3Δ1 leu2Δ0 ura3Δ0 arg4Δ ubp8Δ::KANMX ubp10Δ::NATMX* | 41 |
| YMC309 | *MATa his3Δ1 leu2Δ0 ura3Δ0 arg4Δ ubp8Δ::KANMX ubp10Δ::NATMX rad6Δ::URA3* | 53 |
| YMC336 | *MATa his3Δ1 leu2Δ0 ura3Δ0 arg4Δ ubp10Δ::NATMX rad6Δ::URA3* | 53 |
| YMC337 | *MATa his3Δ1 leu2Δ0 ura3Δ0 arg4Δ ubp8Δ::KANMX ubp10Δ::NATMX bre1Δ::KANMX* | This study |
| YMC339 | *MATa his3Δ1 leu2Δ0 ura3Δ0 arg4Δ ubp8Δ::KANMX ubp10Δ::NATMX lge1Δ::KANMX* | This study |
| YMC27 | *Matα ura3-52, leu2-3,112 his3 trp1Δ LGE1-6HA::TRP1* | This study |
| YMC28 | *Matα ura3-52, leu2-3,112 his3 trp1Δ LGE1-6HA::TRP1 bre1Δ::HIS3* | This study |
| YMC428 | *Matα ura3-52, leu2-3,112 his3 trp1Δ TEL-VII-URA3 lge1Δ::KanMX6* | This study |
| YMC429 | *Matα ura3-52, leu2-3,112 his3 trp1Δ LGE1-6HA::TRP1 rad6Δ::URA3* | This study |

### Spotting assays

The telomeric silencing reporter strain YZS377^53^was transformed with either vector (*LEU2*, CEN) alone or pMC7 derivatives containing either WT *RAD6* or its various mutants. Likewise, telomeric silencing reporter strain YMC428 was transformed with either vector pRS305 (*LEU2*, CEN)^49^or plasmid constructs to express either WT *LGE1* or its mutants. These strains were grown overnight at 30°C with constant shaking in liquid SD media lacking leucine (-LEU). Cells (1 OD_600_ or 1 X 10 ^7^) were harvested. A 10-fold serial dilution was performed, and aliquots were spotted onto solid -LEU media. For the silencing assay, the media additionally contained 5-fluroorotic acid (5-FOA). After spotting, cells were grown at 30°C for 2-3 days before imaging.

### Immunoblotting

Yeast cell extracts were prepared using the TCA lysis method essentially as described previously^41,56^. Log-phase yeast cells (20-25 x 10^7^) were harvested and washed once with phosphate-buffered saline (PBS) and once with 5% tricholoroacetic acid (TCA, Sigma) prior to storing at -80°C. The frozen cell pellets were thawed in 20% TCA and lysed by bead beating. After centrifugation (3000 rpm, 5 min at 4°C), the pellet was resuspended by vortexing in 1X Laemmli buffer (62.5 mM Tris.HCl, pH 6.8, 10% glycerol, 2% SDS, 0.002% bromophenol blue, 2.5% β-mercaptoethanol) and neutralized by adding 2M Tris base before boiling for 8 min in a water bath. The denatured lysate was then clarified by centrifugation (13200 rpm, 10 min at 4°C) and protein concentration was determined using the DC™ Protein Assay (Bio-Rad). Either equal amounts or a serial dilution of the lysates from various strains were resolved using SDS-PAGE and transferred onto a polyvinylidene difluoride membrane. Following incubation with an antigen-specific primary rabbit or mouse antibody and corresponding HRP-conjugated secondary antibody, protein signals were detected by chemiluminescence using Pierce™ ECL Plus Western Blotting Substrate (Thermo Scientific) and autoradiography. The following antibodies were used in immunoblotting: anti-Flag M2 (F3165; Sigma), anti-V5 (46-0708; Invitrogen); anti-HA (39628; Active Motif); anti-Pgk1 (459250; Invitrogen), anti-H2B (39237; Active Motif), anti-H3 (ab1791; Abcam), anti-H3K4me1 (39297; Active Motif), anti-H3K4me2 (399141; Active Motif), anti-H3K4me3 (39159; Active Motif), anti-ubiquityl-Histone H2B (Lys120) (D11) XP® (5546; Cell Signaling); anti-mono- and polyubiquitinylated conjugates monoclonal antibody (clone FK2) (HRP conjugate) (BML-PW0150; Enzo Life Sciences), anti-PCNA/Pol30 (ab221196; Abcam). Anti-Bre1 and anti-Rad6 antibodies were raised in rabbits^28,57^. Anti-Rad6 antibody was purified from rabbit serum as described^57^.

### *In vitro* ubiquitination assay

Recombinant yeast Rad6 or Rad6-150 were expressed and purified from bacteria essentially as described previously ^28^. The ubiquitination reaction was performed in ubiquitination buffer (50mM Tris pH8.0, 50mM KCl,50 mM NaCl, 5mM MgCl2, 5 mM ATP) with 0.1 μM recombinant yeast E1 (R&D Systems),, 50 μM recombinant yeast ubiquitin (R&D Systems), 5 μM Rad6 or Rad6-150, and 2 μM substrate recombinant yeast histone H2B and incubated at 30°C for 30 min, 1 h or 2 h. The reactions were stopped by adding 2X Laemmli sample buffer (Bio-Rad) and resolved in a 12% SDS-PAGE prior to immunoblotting with anti-monoubiquitinated and polyubiquitinated protein antibody (clone FK2), anti-yeast H2B antibody, or anti-Rad6 antibody.

## Acknowledgements

We thank Zu-wen Sun for his input in using DUB deletion strains to examine histone ubiquitination. This work was supported by funds provided by the Department of Radiation Oncology, Huntsman Cancer Institute and NIGMS R01GM127783 to M.B.C.

## Author contributions

Kaitlin S. Radmall: investigation, methodology, formal analysis; Prakash K. Shukla: investigation, methodology, formal analysis, writing -original draft preparation; Mahesh B. Chandrasekharan: conceptualization, investigation, formal analysis, writing -reviewing & editing, supervision, project administration, funding acquisition.

## Availability of Data and Materials

The data supporting the findings of this study are available within the paper and in the supplementary information file. Yeast strains and plasmids are available from the corresponding author, Mahesh B. Chandrasekharan, upon request.

