## supplemental figures for "Structure-function analysis of histone H2B and PCNA ubiquitination dynamics using deubiquitinase-deficient strains"

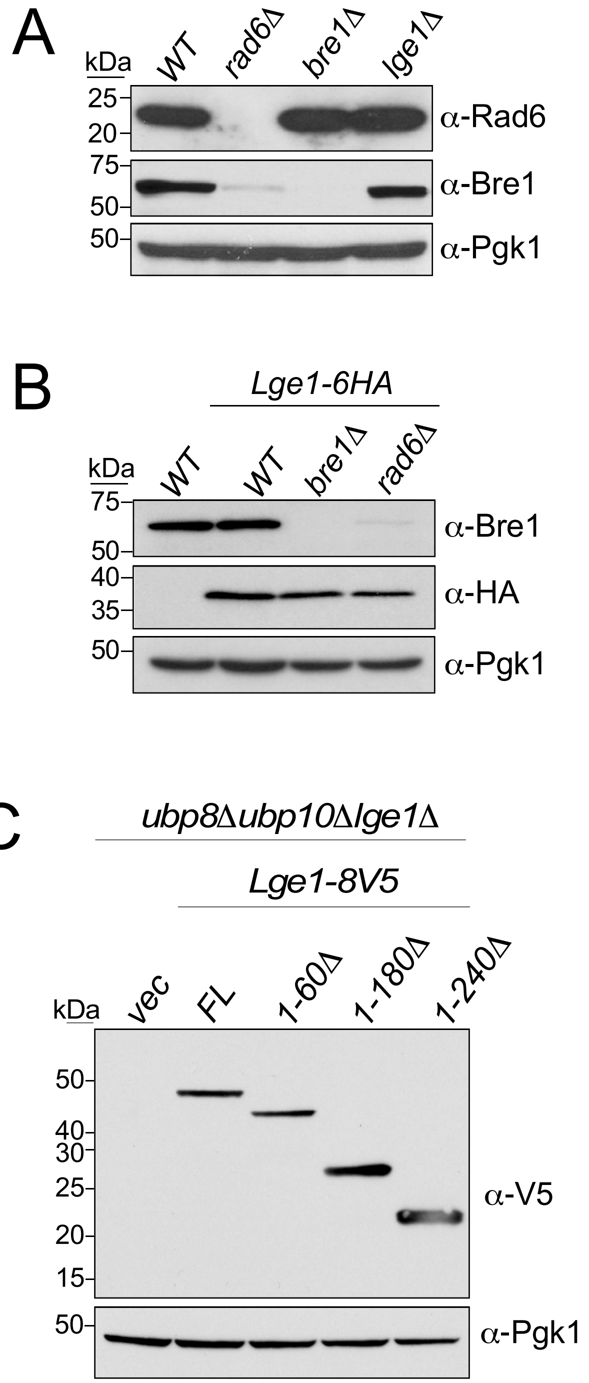


**Figure S1. (A)** Immunoblots for Rad6 and Bre1 in extracts prepared from WT and indicated null mutant strains. **(B)** Immunoblots for Bre1 and HA epitope-tagged Lge1 in extracts prepared from WT and indicated null mutant strains. Extract from a WT strain without HA-tagged Lge1 served as a negative control. **(C)** Immunoblots for V5 epitope-tagged Lge1 or its N-terminal truncation mutants expressed in *ubp8Δubp10Δlge1Δ* null mutant strain. Extract from a strain transformed with empty vector (vec) served as a negative control. Pgk1 levels served as loading control.

```
P06106|Rad6_Scer MSTPARRRLMRDFKRMKEDAPPGVSASPLPDNVMVWNAMIIGPADTPYEDGTFRLLLEFD 60
P23566|Rhp6_Spom MSTTARRRLMRDFKRMQQDPPAGVSASPVSDNVMLWNAVIIGPADTPFEDGTFKLVLSFD 60
P49459|UBE2A_Hsap MSTPARRRLMRDFKRLQEDPPAGVSGAPSENNIMVWNAVIFGPEGTPFEDGTFKLTIEFT 60
P63146|UBE2B_Hsap MSTPARRRLMRDFKRLQEDPPVGVSGAPSENNIMQWNAVIFGPEGTPFEDGTFKLVIEFS 60
                  *** ***********:::* * ***.:*  :*:* ***:*:** .**:*****:* :.*

P06106|Rad6_Scer EEYPNKPPHVKFLSEMFHPNVYANGEICLDILQNRWTPTYDVASILTSIQSLFNDPNPAS 120
P23566|Rhp6_Spom EQYPNKPPLVKFVSTMFHPNVYANGELCLDILQNRWSPTYDVAAILTSIQSLLNDPNNAS 120
P49459|UBE2A_Hsap EEYPNKPPTVRFVSKMFHPNVYADGSICLDILQNRWSPTYDVSSILTSIQSLLDEPNPNS 120
P63146|UBE2B_Hsap EEYPNKPPTVRFLSKMFHPNVYADGSICLDILQNRWSPTYDVSSILTSIQSLLDEPNPNS 120
                  *:****** *:*:* ********:*.:*********:*****::********:::**  *
                                             149 153
                                              |   |
P06106|Rad6_Scer PANVEAATLFKDHKSQYVKRVKETVEKSWEDDMDDMDDDDDDDDDDDDDEAD 172
P23566|Rhp6_Spom PANAEAAQLHRENKKEYVRRVRKTVEDSWES--------------------- 151
P49459|UBE2A_Hsap PANSQAAQLYQENKREYEKRVSAIVEQSWRDC-------------------- 152
P63146|UBE2B_Hsap PANSQAAQLYQENKREYEKRVSAIVEQSWNDS-------------------- 152
                  *** :** *.:::* :* :**   **.**..
```

**Figure S2.** Sequence alignment performed using Clustal Omega for Rad6 and its homologs. Conserved residues (★) and residues with strong (:) or weak similarity (•) are indicated. Uniprot IDs are indicated. Scer, *Saccharomyces cerevisiae*; Spom, *Schizosaccharomyces pombe*; Hsap, *Homo sapiens*.

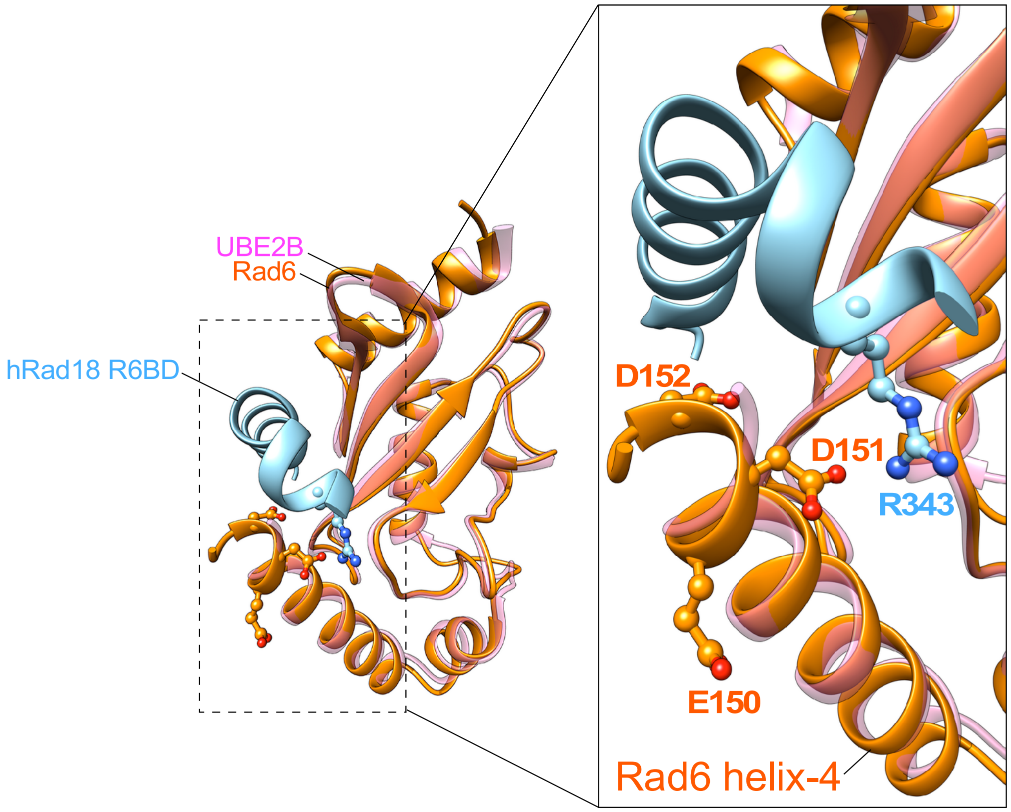


**Figure S3.** Structure of yeast Rad6 (PDB 1AYZ) superimposed on human Rad6b/UBE2B in complex with Rad18 R6BD (i.e., Rad6 binding domain) (PDB 2YBF). Zoomed image shows the helix-4 terminal negatively charged residues of Rad6 with aspartate 151 of Rad6 in close proximity to arginine 343 of the Rad18 R6BD.

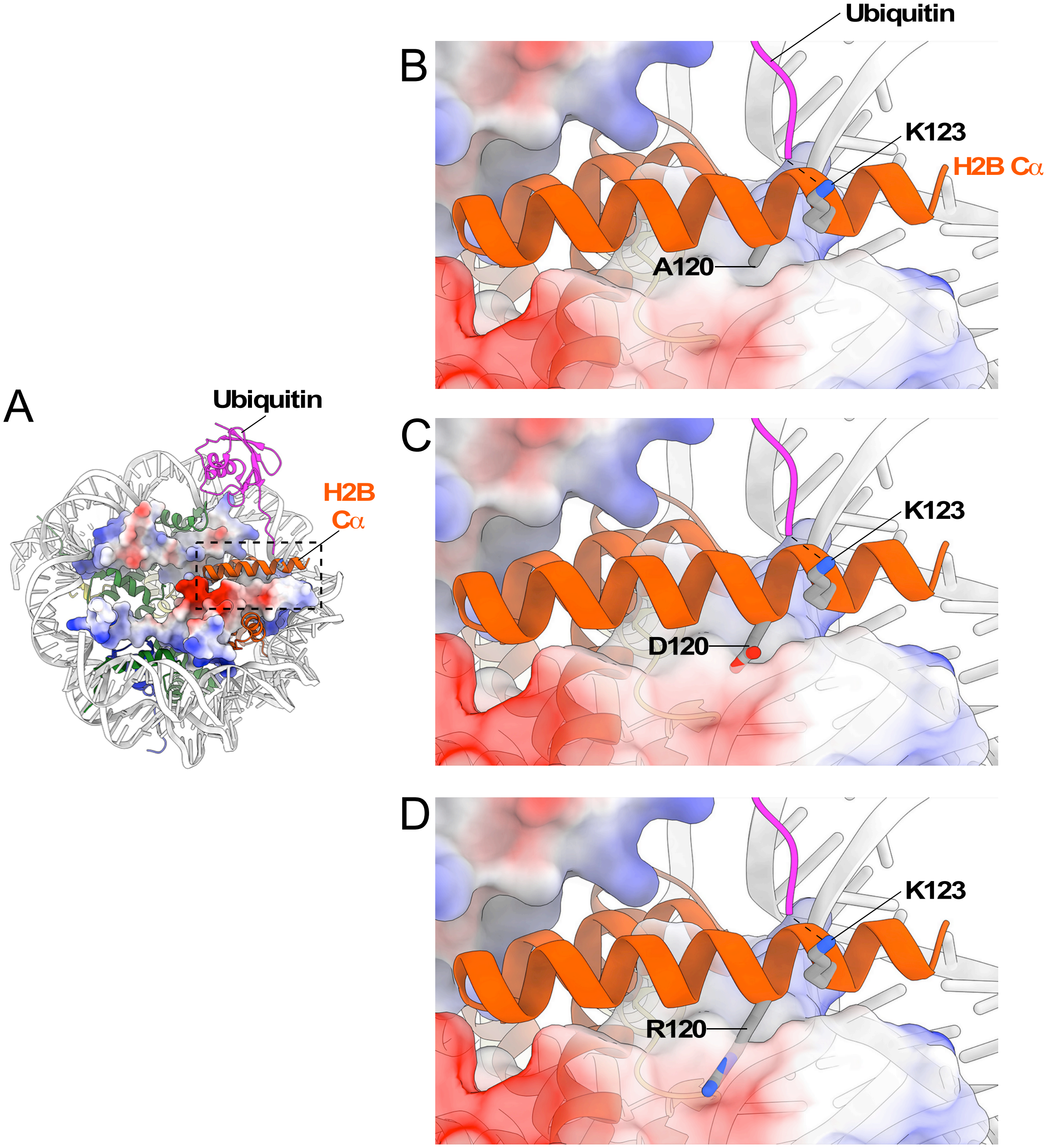


**Figure S4. (A)** Structure of yeast nucleosome (PDB 1ID3) superimposed on the H2BK120ub1 *Xenopus* nucleosome (PDB 4ZUX). The structure for ubiquitin (pink) was retained to obtain a model for the yeast nucleosome with histone H2B (orange ribbon) monoubiquitinated at K123. Histone H2A and H4 are shown in surface charge representation. Gray, hydrophobic; blue, basic; red, acidic. **(B)** Zoomed image of H2B C-terminal helix (Cα). Histone H2B lysine-123 (K123) and alanine-120 (A120) are shown in ball and stick representation. Dashed line indicates isopeptide bond between K123 and glycine at the C-terminal end of ubiquitin. **(C-D)** Models for nucleosomes with C) aspartate and D) arginine substitutions at position 120 in the H2B C-terminal helix. The negatively charged aspartate (D120) resides in a repulsive acidic or hydrophobic environment likely resulting in a conformational change in the H2B C-terminal helix. On the other hand, the positively charged arginine likely interacts with the neighboring acidic residues to stabilize the helix.
